# Oxoglutarate dehydrogenase coordinates myofibril growth by maintaining amino acid homeostasis

**DOI:** 10.1101/2021.12.13.472149

**Authors:** Nicanor González Morales, Océane Marescal, Szilárd Szikora, Miklos Erdelyi, Péter Bíró, Tuana Mesquita, József Mihály, Frieder Schöck

## Abstract

Myofibrils are long intracellular cables specific to muscles, composed mainly of actin and myosin filaments. The actin and myosin filaments are organized into repeated units called sarcomeres, which form the myofibril cables. Muscle contraction is achieved by the simultaneous shortening of sarcomeres and for a highly coordinated contraction to occur all sarcomeres should have the same size. Muscles have evolved a variety of ways to ensure sarcomere homogeneity, one example being the controlled oligomerization of Zasp proteins that sets the diameter of the myofibril. To understand how Zasp proteins effect myofibril growth, we looked for Zasp-binding proteins at the Z-disc. We found that the E1 subunit of the oxoglutarate dehydrogenase complex is recruited to the Z-disc by Zasp52 and is required to sustain myofibril growth. By making specific mutants, we show that its enzymatic activity is important for myofibril growth, and that the other two subunits of the complex are also required for myofibril formation. Using super resolution microscopy, we revealed the overall organization of the complex at the Z-disc. Then, using metabolomic analysis, we uncovered an amino acid balance defect affecting protein synthesis, that we also confirmed by genetic tools. In summary, we show that Zasp controls the local amino acid pool responsible for myofibril growth by recruiting the OGDH complex to the Z-disc.

## Introduction

Striated muscles are long contractile cables that bridge two rigid structures such as bones or regions of the exoskeleton. Muscle contraction is responsible for providing the energy required to displace those rigid units and thus animal movement. Muscles are formed by long intracellular cables, called myofibrils which bridge the ends of muscles, and the coordinated shortening of these myofibrils cause muscle contraction. Myofibrils are themselves composed of tandemly repeated units called sarcomeres. Sarcomeres are built up from a complex array of antiparallel actin and myosin filaments, where actin filaments are anchored to the flanks of the sarcomere at a protein complex called the Z-disc, while the myosin filaments are anchored at the center of the sarcomere at another protein complex called the M-line. Coordinated shortening of the sarcomeres produces myofibril contraction (Gunage, Dhanyasi, Reichert, & VijayRaghavan, 2017; Lemke & Schnorrer, 2017; Luis & Schnorrer, 2021; Nikonova, Kao, & Spletter, 2020).

Because all sarcomeres contract in synchrony, their sizes are identical, and muscles use a variety of ways to ensure sarcomere size homogeneity. This is particularly true for the Indirect Flight Muscle (IFM), a special muscle that evolved in insect lineages to sustain high frequency contractions for prolonged periods of time. One of the adaptations of the IFM are very regular sarcomeres with very small contractile range. In *Drosophila*, the IFM develops during the early pupal stages and then it rapidly grows to fill most of the thoracic space during the late pupal stages (Reedy & Beall, 1993). The muscle growth is a very coordinated process. Myofibrils first form very thin longitudinal cables that stably grow by recruiting cytoplasmic proteins (Katzemich, Liao, Czerniecki, & Schöck, 2013; Loison et al., 2018; Reedy & Beall, 1993; Spletter et al., 2018). The M-line and Z-disc grow together with the myofibril, actively mediating myofibril growth (Gonzalez-Morales, Xiao, et al., 2019; Katzemich et al., 2012; Katzemich et al., 2013; Orfanos et al., 2015).

Local and global mechanisms control muscle growth and sarcomere homogeneity. Local mechanisms acting on individual myofibrils or sarcomeres control sarcomere length, myofibril width and length of the I-band (the region that contains Z-discs and is devoid of myosin filaments). Sarcomere length is controlled by the length of the connecting protein titin (Tskhovrebova & Trinick, 2017) and the fine regulation of actin and myosin dynamics (Molnar et al., 2014; Shwartz, Dhanyasi, Schejter, & Shilo, 2016). Length of the I-bands is controlled by function of the Lasp proteins (Fernandes & Schöck, 2014). Myofibril width is set in place by controlled oligomerization of Zasp proteins at the Z-disc (Gonzalez-Morales, Xiao, et al., 2019). In contrast to these, global mechanisms act on the whole muscle, containing thousands of sarcomere units. One example of a global mechanism is the muscle growth coordination imposed by the continuous increase in tissue tension. As muscles grow, tension builds because of premature muscle contractions and the increase in tendon and cuticle stiffness (Chu & Hayashi, 2021; Lemke & Schnorrer, 2017; Weitkunat, Brasse, Bausch, & Schnorrer, 2017; Weitkunat, Kaya-Copur, Grill, & Schnorrer, 2014). This continuous increase in tension coordinates the growth and the shape of myofibrils and mitochondria (Avellaneda et al., 2021). Other global mechanisms include growth regulation by the Hippo pathway (Kaya-Copur et al., 2021), the role of the E2F/DP heterodimeric transcription factor (Zappia & Frolov, 2016; Zappia, Rogers, Islam, & Frolov, 2019), and the role of insulin (Demontis & Perrimon, 2009). The Hippo pathway and E2F/DP separately coordinate myofibril and mitochondrial growth rate by promoting the expression of myofibril and mitochondria proteins (Kaya-Copur et al., 2021; Zappia et al., 2019). Despite all these well described mechanisms, the interconnections between the global growth cues and the local growth mechanisms have yet to be deciphered.

To gain insights into the mechanisms of Zasp proteins promoting myofibril growth, we screened for Z-disc proteins recruited by Zasp involved in myofibril diameter size regulation. Zasp proteins are members of the ALP/Enigma family, they consist of a PDZ, a ZM, and up to-4 C-terminal LIM domains. They localize to the Z-disc, with their LIM domains at the very center of the disc, and their PDZ domain at the periphery (Katzemich et al., 2013; Szikora et al., 2020). Through the PDZ domain, Zasp binds actinin and establishes the structural core of the Z-disc (Liao et al., 2016). Actinin anchors actin filaments from opposing sarcomeres. Zasp exists in two forms, a blocking and a growing form, with opposite roles during myofibril diameter growth (Gonzalez-Morales, Marsh, et al., 2019; Gonzalez-Morales, Xiao, et al., 2019; Katzemich et al., 2013). The blocking forms prevent recruitment of the growing isoforms to the Z-disc, whereas the growing isoforms recruit Zasp proteins to the Z-disc. The self-association of Zasp is mediated by a physical interaction between its LIM domains and ZM domain (Gonzalez-Morales, Xiao, et al., 2019).

We report that 2-Oxoglutarate Dehydrogenase (OGDH/E1), a crucial enzyme in the tricarboxylic acid (TCA) cycle, is recognized by the LIM domains of Zasp. OGDH/E1 is recruited to the growing Z-disc, and its function is required for myofibril growth. The TCA cycle is a loop of chemical reactions and constitutes a metabolic buffering system. Anaplerotic metabolic pathways replenish the TCA cycle, while cataplerotic reactions use the TCA cycle metabolites (Martinez-Reyes & Chandel, 2020; Owen, Kalhan, & Hanson, 2002). An important TCA step is the conversion of 2-oxoglutarate into succinyl-CoA catalyzed by the OGDH complex, a giant enzyme cluster composed of multiples of three subunits (Tretter & Adam-Vizi, 2005). The Dihydrolipoyllysine-residue succinyltransferase subunit (DLST/E2/CG5214) serves as a structural core unit (Skalidis, Tuting, & Kastritis, 2020). The OGDH/E1/Nc73EF and Dihydrolipoyl dehydrogenase (DLD/E3/CG7430) subunits sit in the periphery of the E2 core (Larkin et al., 2021; Skalidis et al., 2020; Tretter & Adam-Vizi, 2005) (Chen et al., 2015; Gruntenko et al., 1998; Yoon et al., 2017). The OGDH/E1 subunit recognizes the 2-oxoglutarate substrate and provides specificity to the complex (Tretter & Adam-Vizi, 2005). Here we show the nanoscale distribution of these subunits in the Z-disc and demonstrate that the absence of OGDH/E1 results in severe alterations in amino acid metabolism of the muscle cells. Together, these data establish a novel link between a local, Zasp-dependent myofibril growth control mechanism and a global myofibril growth control mechanism, amino acid availability.

## Results

Because Zasp proteins are often insoluble, to find proteins recruited by Zasp to the growing Z-disc, we used a bioinformatic approach to look for proteins with similar evolutionary rates as the Zasp proteins. Evolutionary Rate Covariation (ERC) is a measurement of shared evolutionary history between proteins (Clark & Aquadro, 2010). We retrieved the ERC values for Zasp52, Zasp66, and actinin, and matched them to all *Drosophila* proteins with available ERC values (roughly 11,100 proteins). We selected proteins with ERC values greater than 0.5 with at least two of the three bait proteins. Then, we used an automatic clustering method to group the candidate proteins. Two clusters were obtained, the Zasp52/Zasp66 and the Zasp52/actinin groups (**Figure 1A**), possibly reflecting the two roles of Zasp52: stabilize actinin and recruit other Zasp proteins. As we primarily aimed for Z-disc proteins, we asked which of the candidate proteins colocalizes with Zasp and actinin at the Z-disc. To do so, we obtained tagged versions of the candidate proteins and analyzed their intracellular localization. To avoid the signal coming from the mitochondria, we used a glycerinating step to wash away the mitochondria (Xiao, Schöck, & González-Morales, 2017). Only TER94/VCP and Nc73EF/OGDH showed a clear Z-disc localization (**Figure 1B**). TER94/VCP is an AAA+ ATPase linked to protein quality control with known muscle functions (Chang et al., 2011; Zhang, Mishra, Hay, Chan, & Guo, 2017). Oxoglutarate dehydrogenase (OGDH) is a critical enzyme in the TCA cycle (Martinez-Reyes & Chandel, 2020).

**Figure 1.**
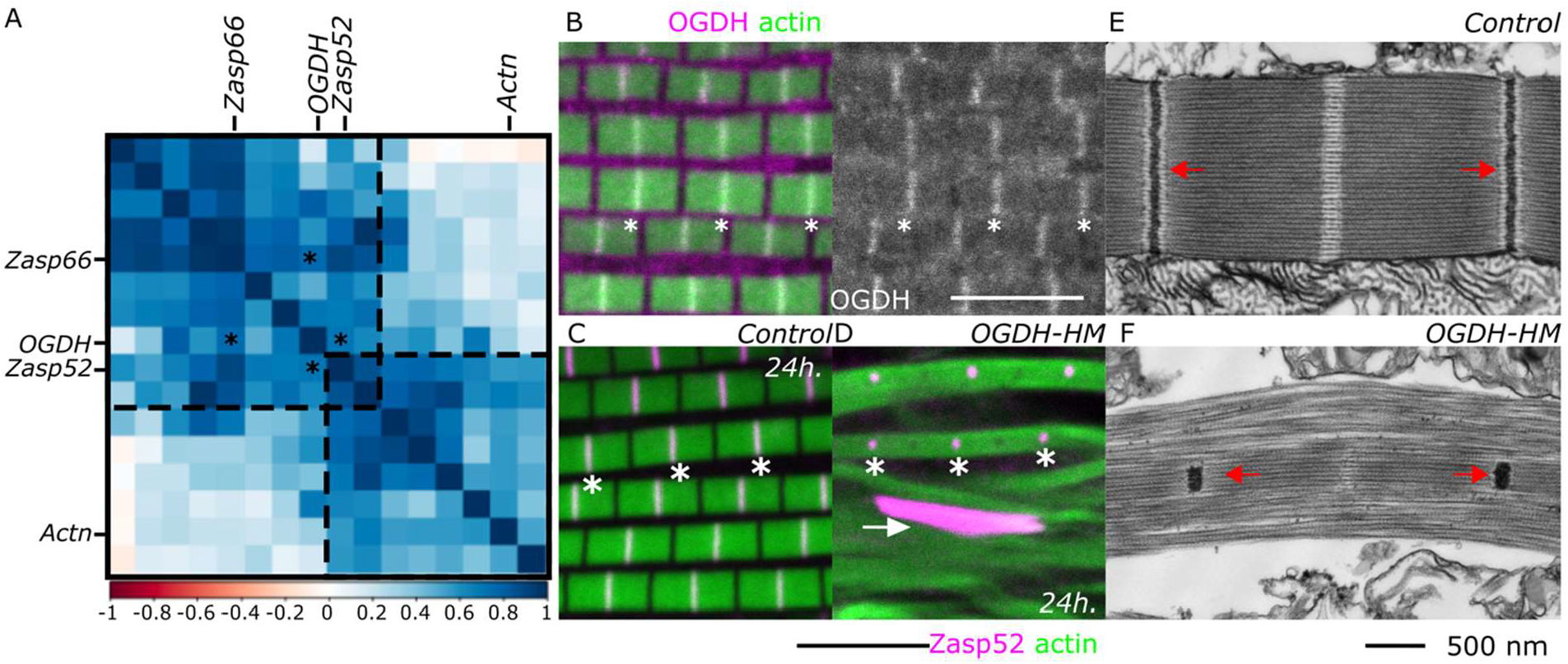
OGDH localizes to the Z-disc and is required for myofibril growth. (A) Heatmap showing the ERC values from all the hits in the ERC screen ordered by automatic clustering. Note the Zasp52/Zasp66 and the Zasp52/actinin clusters. The color scale of the ERC values from -1 to +1 is at the bottom. The positions of Actn, Zasp52, Zasp66, and OGDH are indicated. (B) Confocal image of OGDH[MI06026-GFSTF.1] muscles, OGDH-GFP localizes to the Z-disc. Composite and single-channel images are shown. (C-D) Confocal images of control (C) and OGDH-HM (D) muscles with Zasp52-mCherry as a Z-disc marker (magenta) and actin filaments as reference (green). In OGDH-HM muscles, the Z-discs do not grow to their final size and appear small, protein aggregates are occasionally present. (E-F) TEM images of control and OGDH-HM sarcomeres. The Z-discs are the black electron-dense structures indicated by red arrows. The M-line is the white region at the center of the sarcomere. In the control image, the Z-disc is well defined and spans the whole diameter of the sarcomere. In OGDH-HM muscles, the Z-disc is reduced to a tiny clump at the middle of the myofibril. White asterisks show selected Z-discs in B-D. Scale bars are 5 µm in B to D and 500 nm in E and F.

Because OGDH localization at the Z-disc was unexpected, we decided to further study its role in myofibril assembly. First, we analyzed the consequences of inactivating OGDH. We removed OGDH from the developing indirect flight muscles using Act88f-Gal4 in combination with an RNAi directed against all *OGDH* isoforms (HMS00554, we refer to these flies as OGDH-HM). We noted that all the OGDH-HM flies were flightless, and all their myofibrils were abnormal as compared to controls (**Figure 1C and D**). OGDH-HM muscles had very small Z-discs and occasionally protein aggregates were visible in the cytoplasm (**Figure 1C and D**). Next, we analyzed the localization of other myofibril proteins in OGDH-HM muscles. Zasp66 and actinin localized to both the Z-discs and the aggregates, Sls/titin only to the Z-disc, and obscurin/unc-89 was restricted to the M-line (**Figure 1 – figure supplement 1)**. Also, we used transmission electron microscopy to characterize the phenotype in more details, and we noted that in OGDH-HM flies the Z-discs, located at the center of the fibrils, were severely reduced in size (**Figure 1E-F**), suggesting a Z-disc growth defect.

To test the specificity of the RNAi knock down, we used other approaches to reduce the function of OGDH in the muscles. We used two other RNAi lines that target different sequences of the *OGDH gene* (GD12778 and GD50393) and an indirect flight muscle specific CRISPR-Cas9 based method targeting the OGDH gene (OGDH-CRISPR, TKO.GS00550). All these conditions had muscles with smaller Z-discs and protein aggregates (**Figure 1 – figure supplement 1)**, confirming the OGDH requirement for proper myofibril diameter growth. The Z-discs were smaller in case of OGDH-HM and OGDH-CRISPR than in the two other RNAi conditions, whereas the aggregates were most common in OGDH-CRISP, followed by OGDH-HM, GD50393, and GD12778 (**Figure 1 – figure supplement 1**). Thus, overall, all the four methods resulted in similar muscle defects, and the slight differences observed are likely to reflect the effectiveness of reducing OGDH levels in the different conditions.

After the initial phenotypic characterization of OGDH, we explored the physical association between Zasp52 and OGDH proteins. First, we used a yeast two-hybrid assay to test for protein-protein binding. Yeast expressing Zasp52 and OGDH grow on selective media, suggesting a physical interaction, while yeast expressing Zasp66 or Actinin together with OGDH did not grow (**Figure 2 and data not shown**). We then used this assay to find the binding site mediating the interaction between OGDH and Zasp52. First, we paired OGDH with all the possible individual domains of Zasp52. Only Zasp52-LIM2a and LIM2b together with OGDH restored yeast growth (**Figure 2A)**. We have previously shown that the LIM domains of Zasp52 bind to the ZM domains (Gonzalez-Morales, Xiao, et al., 2019). We noted a region in OGDH that weakly aligns to the ZM domain of Zasp66 and asked if this sequence would mediate OGDH-Zasp52 binding as well (**Figure 2B, C**). We made a mutant OGDH version (referred to as OGDH-BM, short for <u>B</u>inding <u>M</u>utant) that lacks this sequence, and we found that Zasp52 is unable to bind OGDH-BM (**Figure 2D**). The LIM2a or LIM2b domains are also unable to bind OGDH-BM in Y2H assays (**Figure2E**).

**Figure 2.**
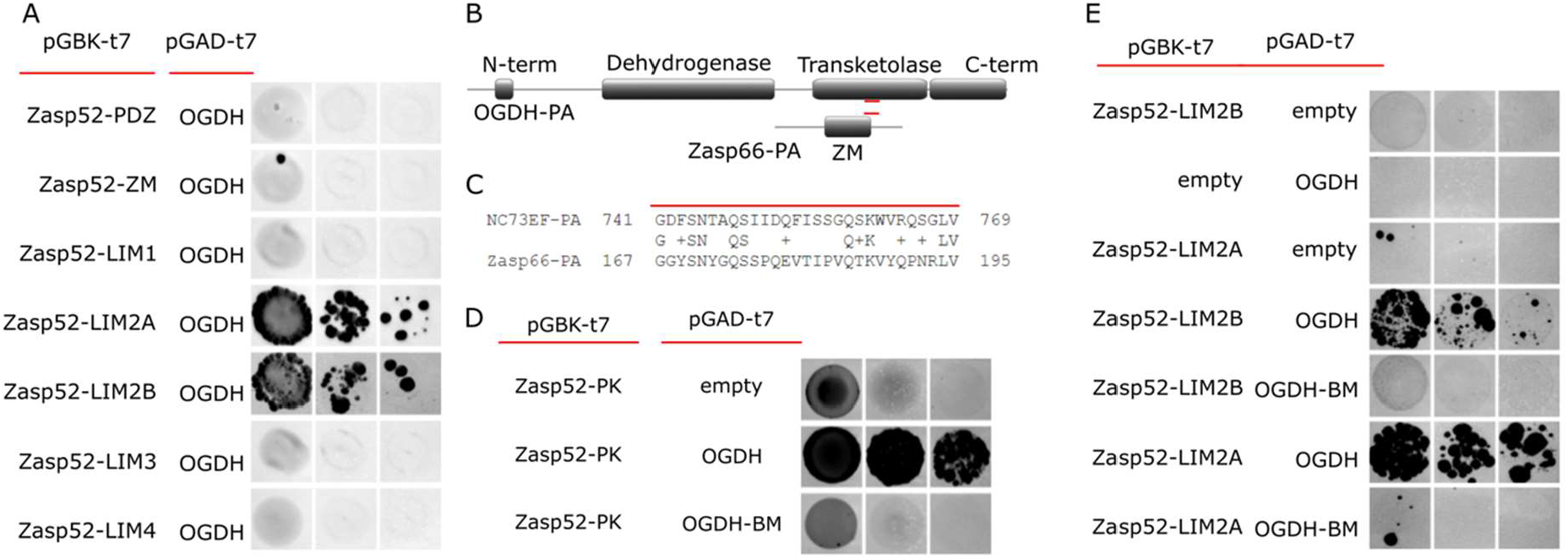
The LIM2 domain of Zasp52 binds OGDH through a ZM-like sequence. (A) Y2H assays showing LIM2a and LIM2b are the only domains in Zasp52 that bind OGDH. (B) Cartoon of OGDH-PA and Zasp66-PA proteins with protein domains highlighted. The short sequence common to both proteins is marked by a red line. OGDH-BM has this short sequence removed. (C) Alignment of the common sequence between OGDH-PA and Zasp66-PA isoforms. (D) Y2H assays showing that Zasp52-PK binds to OGDH but the interaction is lost in OGDH-BM, a mutant version that lacks the common short sequence. (E) Y2H assays showing LIM2a and LIM2b bind the short sequence in OGDH. Serial dilutions are shown from left to right in A, D, and E (OD: 0.1, OD: 0.01, and OD: 0.001).

Because both forms localize to the Z-disc when overexpressed, we used Bimolecular Fluorescence complementation assays to test Zasp/OGDH direct interaction at the Z-disc (Marescal, Schöck, & Gonzalez-Morales, 2020). Muscles expressing OGDH fused to the N-terminal portion of YFP and Zasp52 fused to the complementary C-terminal portion of YFP show a clear fluorescent signal at the Z-disc (**Figure 3A**), suggesting OGDH and Zasp bind at the Z-disc. Importantly, the signal is lost in OGDH-BM conditions (**Figure 3B and C**). We then tested the role of Zasp52 in recruiting OGDH to the Z-disc. We imaged OGDH-GFP in two *Zasp52* mutants. *Zasp52*^*MI02988*^ affects the PDZ domain but not the LIM domains and *Zasp52*^*MI00979*^ forces a stop before the last three LIM domains but leaves the PDZ domain intact. OGDH localizes to the Z-disc in control and in *Zasp52*^*MI02988*^ mutant muscles but not in *Zasp52*^*MI00979*^ mutants (**Figure 3D-F**), further supporting the involvement of the LIM domains in OGDH binding.

**Figure 3.**
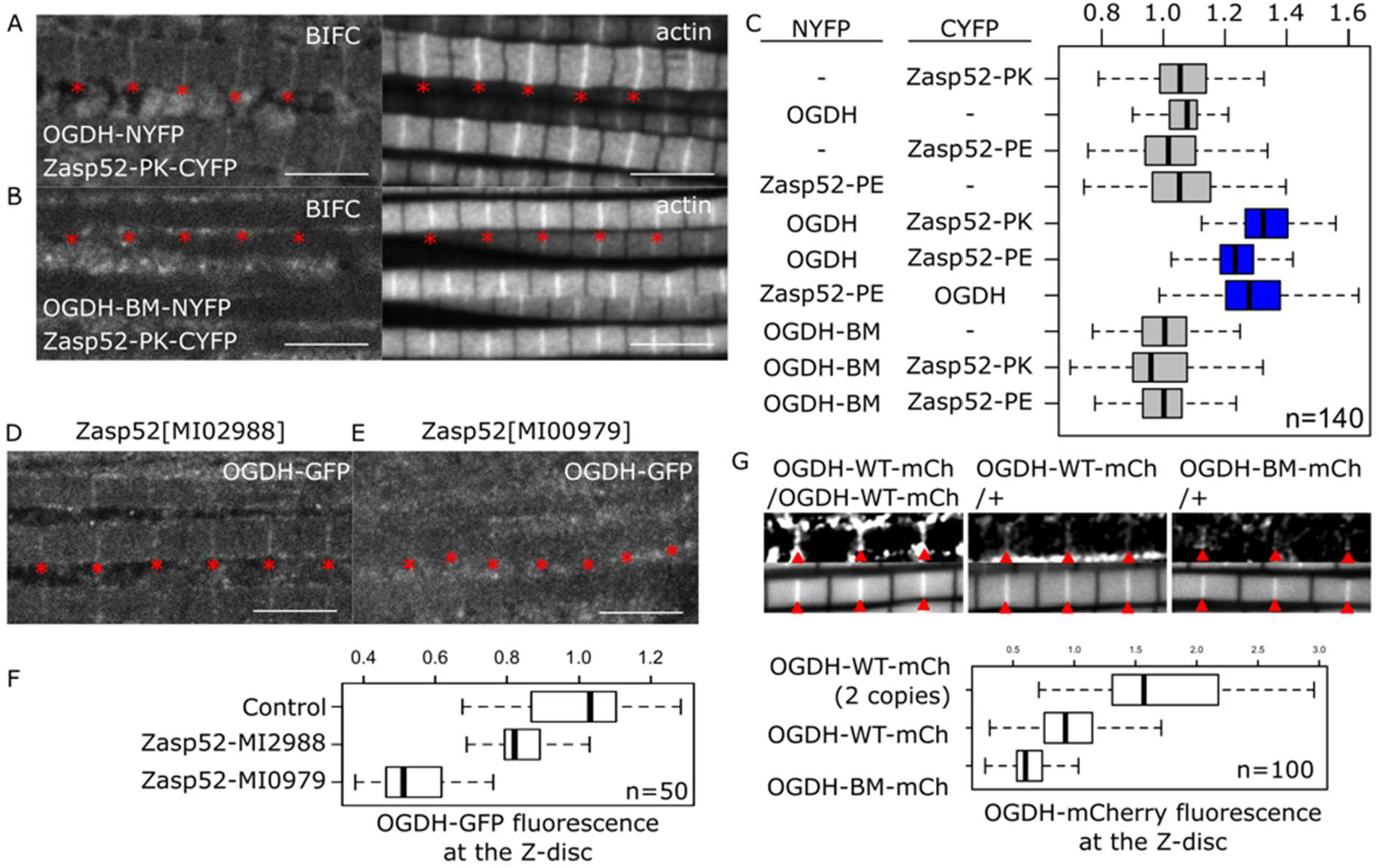
Zasp52 recruits OGDH to the Z-disc. (A, B) Confocal images of muscles showing BiFC fluorescence and actin staining for reference. (A) Zasp52-PK and OGDH form a complex at the Z-disc, BiFC signal is detected at the Z-disc. (B) OGDH-BM does not bind Zasp52-PK at the Z-disc, BiFC signal is not detected at the Z-disc. (C) Plots of the BiFC fluorescence intensity values relative to background noise. Physical interaction is detected between Zasp52 and OGDH, colored in blue. (D, E) Confocal images showing the localization of OGDH-GFP in two *Zasp52* mutants. (D) In *Zasp52*^*MI02988*^ mutants, OGDH is properly localized to the Z-disc. (E) In *Zasp52*^*MI00979*^ mutants, OGDH-GFP is not detected at the Z-disc. (F) Boxplot of OGDH-GFP intensities in control and *Zasp52* mutant backgrounds. (G) Confocal images of wild type or binding mutant OGDH forms tagged with mCherry and box plot of the fluorescence intensity. Flies with one or two copies of OGDH-wt-mCh show Z-disc localization. The OGDH-BM-mCh form has reduced Z-disc localization compared to the wild type. Red arrows denote the Z-discs. The scale bars in all panels are 5 µm.

Because overexpression studies are prone to subcellular localization artifacts, we analyzed the endogenous OGDH protein as well. To this end, we created OGDH mutant alleles by incorporating an mCherry tag either into the wild type OGDH or the OGDH-BM form (*OGDH-WT-mCh* and *OGDH-BM-mCh*, respectively). Homozygous *OGDH-WT-mCh* flies are viable and their myofibrils develop properly. In contrast, homozygous *OGDH-BM-mCh* are lethal. As heterozygotes, both alleles develop normal myofibrils. The Z-disc localization of OGDH-BM-mCh is diminished roughly by half compared to the OGDH-WT-mCh control (**Figure 3G**), suggesting that Zasp binding of OGDH is, at least partially, required for Z-disc localization.

Next, we wanted to address whether the enzymatic activity of OGDH is important for Z-disc formation.We made an enzymatic dead mutant of the endogenous *OGDH* by changing two residues known to mediate substrate binding (Frank, Price, Northrop, Perham, & Luisi, 2007). We compared the phenotype of the OGDH enzyme dead mutant (*OGDH*^*ED*^) to that of *OGDH*^*T2A*^, an M[Trojan-Gal4] insertion that introduces premature stop codons in all OGDH isoforms. Because the homozygous OGDH^*ED*^ and *OGDH*^*T2A*^ mutants die as embryos, we imaged the body wall muscles in late stage 17 embryos of both mutants using Zasp52-GFP as a Z-disc marker (**Figure 4A-C**). In the controls, the Z-discs are equidistant and have a regular appearance (**Figure 4A**). In *OGDH*^*T2A*^ mutants the Z-disc arrangement is completely lost (**Figure 4B)**. In OGDH^*ED*^ mutants, the Z-disc arrangement is severely affected, such as in OGDH^*T2A*^ (**Figure 4C)**. Because OGDH normally functions as one of the three subunits of the OGDH protein complex, we used tissue specific CRISPR-Cas9 targeted mutagenesis to examine the function of the three OGDH complex subunits. We expressed Cas9 together with gRNAs targeting *OGDH/E1, DLSTE2, or DLD/E3* in the indirect flight muscles and observed a very similar sarcomere phenotype in all three cases, characterized by a strong impairment of myofibrillar organization and sarcomere structure (**Figure 4D-G**). Collectively, these data strongly argue that enzymatic function of the OGDH complex is crucial for proper sarcomere arrangement, including Z-disc organization.

**Figure 4.**
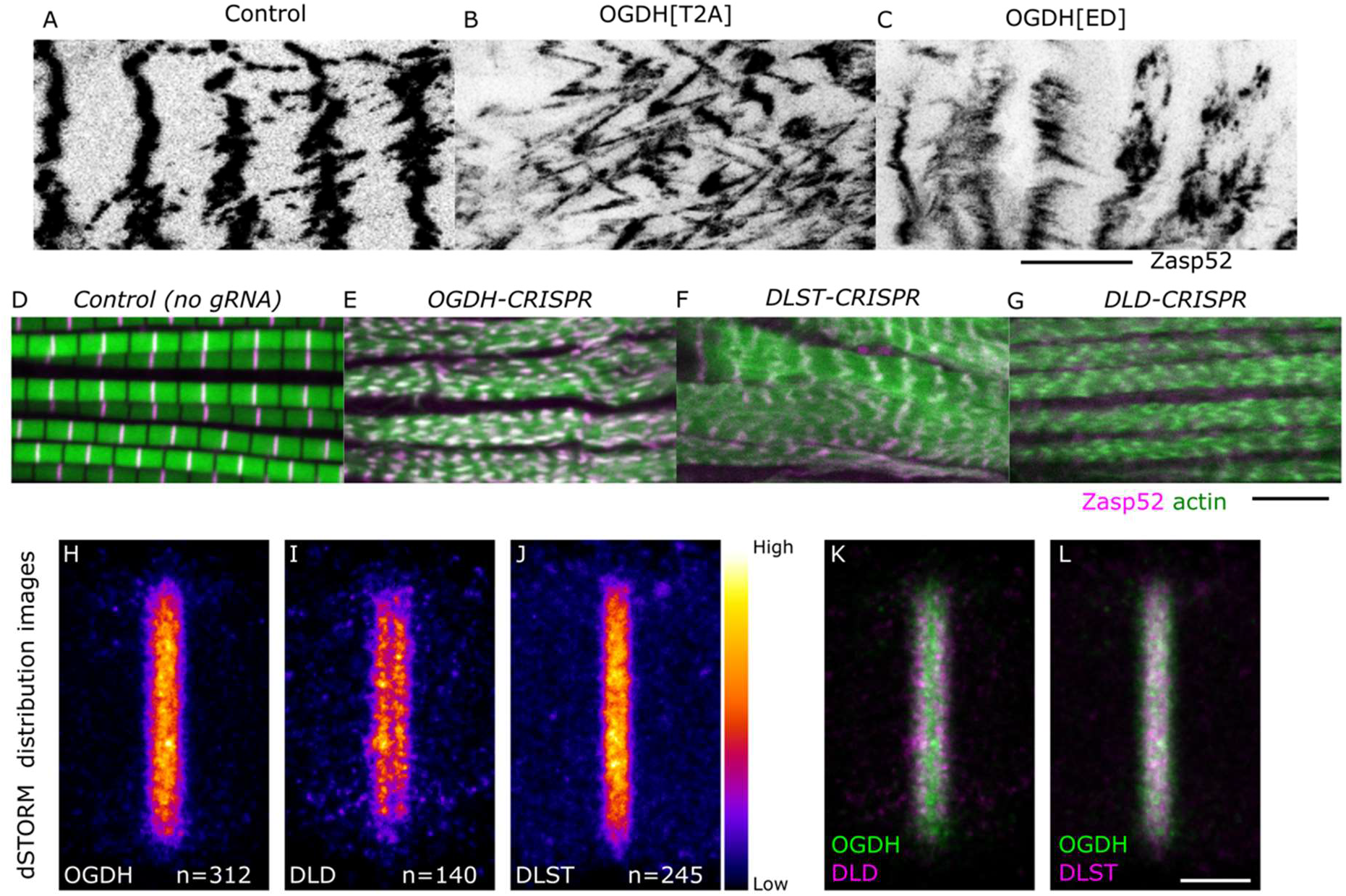
The OGDH complex at the Z-disc. (A-C) Confocal images of Zasp52-GFP in embryonic body wall muscles showing the muscle defects associated with OGDH mutants. (A) In control muscles, the Z-discs, marked by Zasp52, are homogeneous and equidistant. (B) In OGDH^*T2A*^ mutants the stereotypic Z-disc pattern is mostly gone. (C) In OGDH^*ED*^ mutant the Z-disc arrangement is severely affected. (D-G) Confocal images of muscles with CRISPR-induced mutations of the OGDH complex subunits. The E1 subunit OGDH (E), the E2 subunit DLST (F), and the E3 subunit DLD (G). Mutations in any of the OGDH complex subunits result in very thin myofibrils that tend to stack on each other. Zasp52 marks the Z-disc in magenta and actin filaments are shown in green. (H-L) Super resolution dSTORM images subunits of the OGDH complex as intensity heat maps. (E) The E1 subunit, OGDH, localize to a wide band at the Z-disc core. (F, H) The E3 subunit, localizes to two discrete bands that overlaps the periphery of the E1 subunit distribution. (G, I) The E2 core subunit has a distribution pattern like the E1 subunit. Scale bar in A-C is 10 µm. The scale bar in D-G is 5 µm. The Scale bar in E-I is 500 nm.

To further test the requirement of all three subunits, we used super resolution microscopy to determine the precise localization of the subunits by employing a previously validated dSTORM (direct Stochastic Optical Reconstruction Microscopy) method (Szikora et al., 2020). We found that the distribution of DLST, the core subunit, and that of the OGDH molecules are largely overlapping with each other at the Z-disc **(Figure 4J and L)**. The third subunit, DLD, also accumulates at the Z-disc, but it localizes into two discrete bands alongside the Z-disc **(Figure 4I and K**). The double-band pattern of DLD still exhibits a significant overlap with that of DLST and OGDH, therefore these super resolution data suggest that two OGDH complexes assemble at either side of the Z-disc center, with the DLD subunits concentrated on the outer periphery **(Figure 4 - figure supplement 1 and 2)**.

Given the critical role of OGDH in the TCA cycle, we next asked if depletion of other TCA cycle components would affect sarcomere assembly. We expressed RNAi transgenes targeting the major components of the TCA cycle (**Figure 5A**), and strikingly, we observed sarcomere phenotypes in 83% of them (**Figure 5B-H, Table 1**). Most notably, Aconitase, Isocitrate Dehydrogenase, OGDH complex E2 subunit and Succinyl CoA-Synthetase, catalyzing sequential steps in the cycle, had the most dramatic effects (**Figure 5B**), often exhibiting a complete disintegration of the myofibril structure (**Figure 5E, F, and H, red arrows**) or myofibrils with more subtle defects (**Figure 5D, F-H, orange arrows**). In case of the few TCA components that do not affect sarcomere structure, enzyme redundancy is a likely explanation for the lack of phenotype. As the TCA cycle fuels the electron transport chain, which generates ATP, we also tested the role of ATP by analyzing muscles with a compromised electron transport chain. Removing Cox5a, an essential subunit of the Cytochrome c Oxidase (Mandal, Guptan, Owusu-Ansah, & Banerjee, 2005), does not result in myofibril defects despite a strong reduction in ATP levels (**Figure 5B, Figure 5 - figure supplement 1)**, indicating that a reduction in ATP levels does not account for the defects observed in the TCA cycle enzymes.

**Figure 5.**
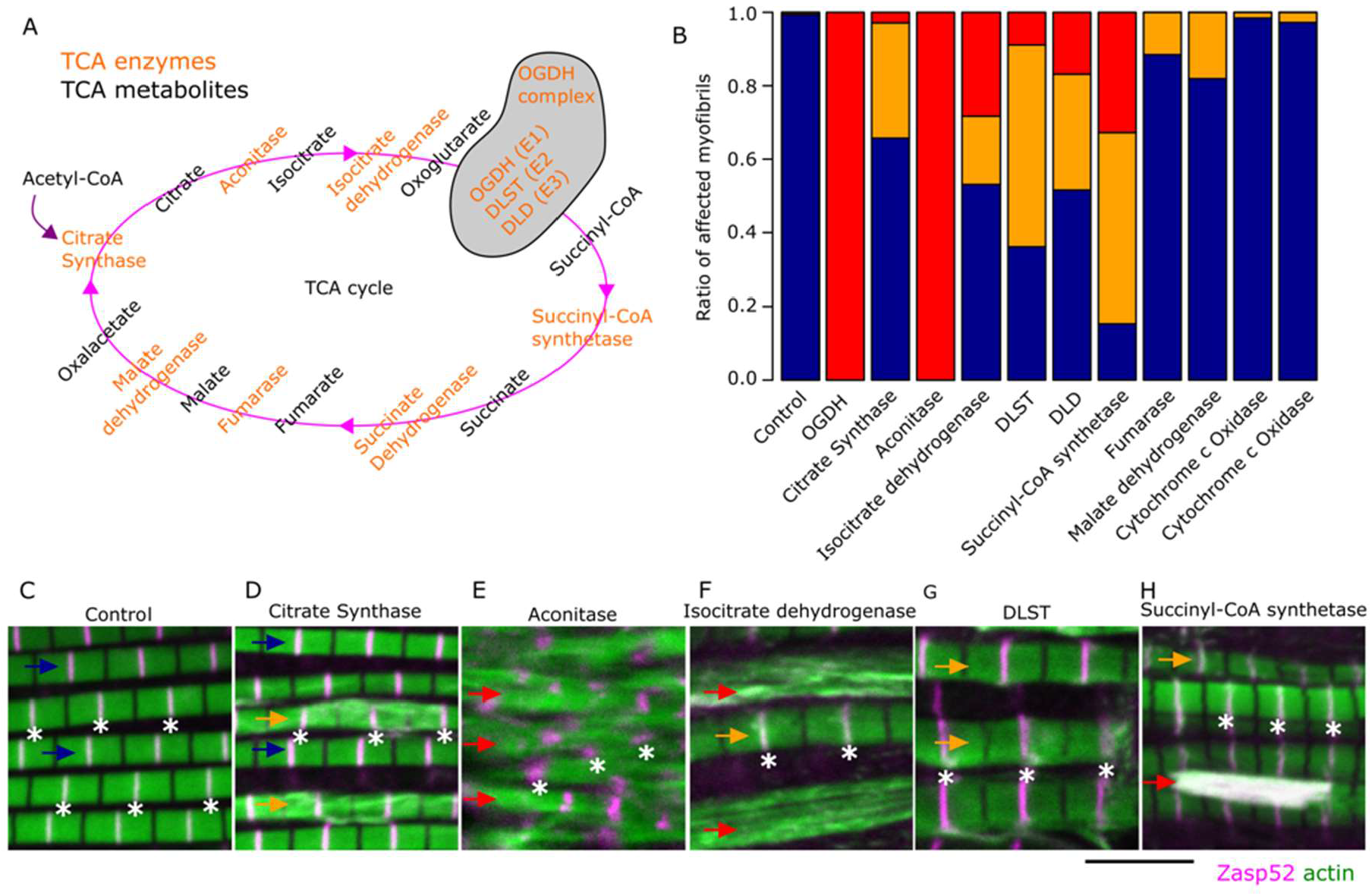
The TCA cycle is involved in sarcomere growth. (A) Cartoon of the TCA cycle in *Drosophila*, the enzyme-encoding genes are orange, and the metabolites are black. The TCA cycle is a loop of chemical reactions and constitutes a metabolic buffering system. The cycle starts when Citrate Synthase combines acetyl-CoA with oxaloacetate. Then, Aconitase converts citrate into isocitrate. Then, there are two oxidative decarboxylation steps. The homodimeric Isocitrate Dehydrogenase converts isocitrate into oxoglutarate. In the second decarboxylation step, the OGDH complex, composed of three subunits (*OGDH/E1, DLST/E2*, and *DLD/E3*), converts oxoglutarate into succinyl-CoA. Next, Succinyl-CoA synthetase converts succinyl-CoA into succinate. Then, Succinate Dehydrogenase oxidizes succinate to fumarate. Succinate dehydrogenase is also part of the electron transport chain. Then, Fumarase converts fumarate to malate. Finally, Malate dehydrogenase reconstitutes the oxaloacetate. (B) Stacked bar plot of the ratio of myofibrils that appear normal (blue), with defects (orange), and without sarcomere structure (red) in different RNAi conditions. (C-H) Confocal images showing the myofibril phenotypes of select TCA cycle enzymes. Examples of myofibril defects are denoted by colored arrows that match the colors used in B. Asterisks mark the position of selected Z-discs (C) The control genotype (*Act88F-Gal4, Zasp52-mCherry*). (D) Depletion of Citrate Synthase causes myofibril streaming (orange arrows). (E) Depletion of Aconitase causes a complete loss of myofibril structure (red arrows). (F) Depletion of Isocitrate Dehydrogenase causes loss of myofibril structure (red arrows). (G) Depletion of DLST/E2 subunit causes strong myofibril defects (orange arrows). (H) Depletion of Succinyl-CoA synthetase causes a loss of myofibril structure (red arrow) and myofibril defects (orange arrow). In C-H, Zasp52 marks the Z-discs in magenta. Actin filaments are shown in green. Scale bar in C-H is 5 µm.

The TCA cycle is a metabolic hub that connects many aspects of cellular homeostasis including the biosynthesis of many amino acids (Martinez-Reyes & Chandel, 2020). To investigate how the loss of OGDH affects muscle metabolism, we performed metabolomic analysis of OGDH-HM and control muscles (**Figure 6A**). Consistent with a role in the TCA cycle, we observed a 2.7-fold increase in oxoglutarate accumulation, indicating the sensitivity of the approach. Interestingly, this change was paralleled with abnormal levels of numerous amino acids. The most extreme cases are Histidine and β-alanine, which are almost absent compared to control muscles (**Figure 6A and B**). Homoserine, and Aspartic acid are also less abundant in OGDH-HM muscles (**Figure 6B**), whereas Valine, Tyrosine, Leucine, Lysine, Isoleucine, Phenylalanine, and Sarcosine, a glycine intermediate, accumulate in OGDH-HM muscles (**Figure 6B**). Overall, these data suggest a widespread effect on amino acid metabolism.

**Figure 6.**
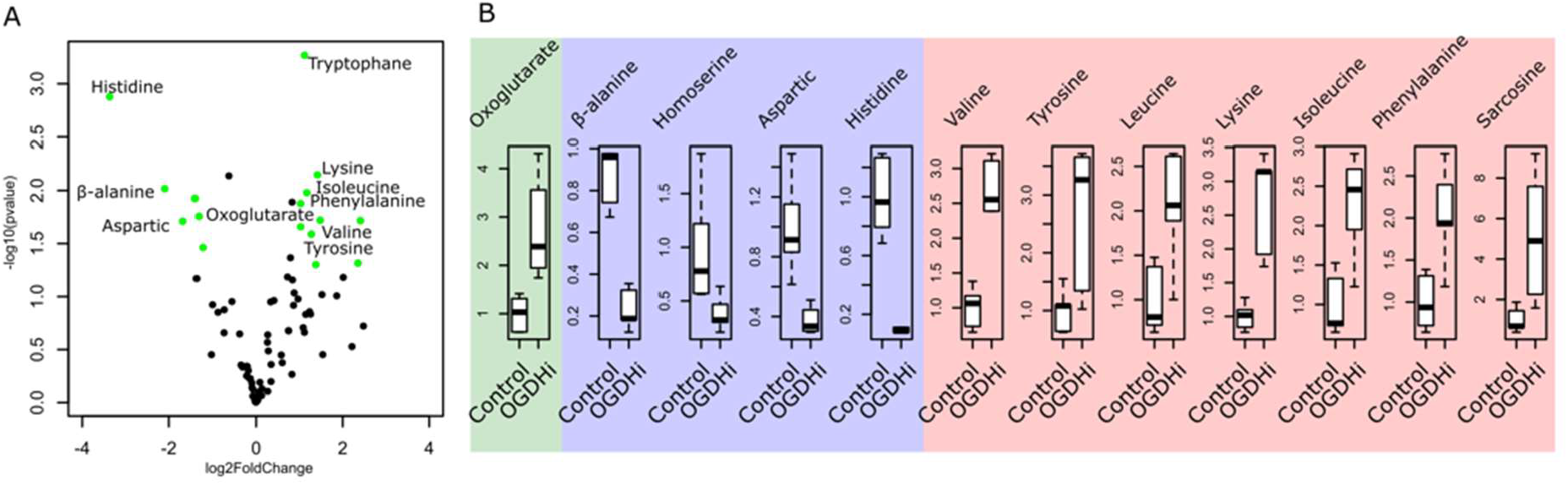
Gas chromatography-mass spectrometry metabolomic analysis of OGDH-HM reveals a defect in amino acid balance. (A) Volcano plot of all the metabolites analyzed in control (Act88F-Gal4, Zasp52-GFP) and OGDH-HM (Act88F-Gal4, UAS-OGDH-HM, Zasp52-GFP) samples. Significance -log10(pvalue) and fold-change log2 are on the y- and x-axes, respectively. Each dot is a unique metabolite, the green dots are metabolites with p values smaller than 0.05 and absolute log2FoldChange values greater than 1. The names of selected metabolites are shown. Note the accumulation of oxoglutarate and the imbalance in many amino acids. (B) Boxplots of selected metabolites in control and OGDH-HM (OGDHi) samples. The green panel shows oxoglutarate is ∼ 3 times more concentrated in OGDH-HM than in controls, validating the approach. The blue panel shows the reduction of β-alanine, homoserine, aspartic acid, and histidine in OGDH-HM compared to the control samples. The red panel shows the accumulation of several amino acids. Sarcosine is an amino acid intermediate of glycine metabolism.

We then hypothesized that the problems in protein synthesis are a likely cause of the myofibril growth defects in OGDH depleted muscles. It is well known that when amino acids are missing, cells actively block protein synthesis by reducing ribosome biogenesis or by actively degrading them (Destefanis, Manara, & Bellosta, 2020; Iadevaia, Liu, & Proud, 2014). To test this possibility, we artificially blocked protein synthesis by expressing the A subunit of the ricin toxin in muscles during different developmental periods (Moffat, Gould, Smith, & O’Kane, 1992). Consistent with our model, ricin expression during early development phenocopies the complete absence of sarcomere structure obtained by inducing CRISPR-Cas9 mutations of any of the subunits of the OGDH complex (**Figure 7A ∼28h**). In contrast, ricin expression slightly after the first appearance and before sarcomere growth, phenocopies the small Z-disc phenotype of RNAi depletion of OGDH (**Figure 7A ∼32 h and ∼56 h**). Finally, ricin expression after the growth period ∼80 h has little effect on sarcomere structure or size (**Figure 7A ∼80 h**). We then asked if the number of ribosomes was affected by OGDH depletion. We used a GFP tagged version of the ribosomal protein RpS5a (Kong et al., 2019). In control muscles, RpS5a localized to the perinuclear region and the Z-discs (**Figure 7B, D**), but in OGDH depleted muscles RpS5a fluorescence was mostly missing (**Figure 7C, E-F**), suggesting ribosome degradation or a stop in ribosome biogenesis.

**Figure 7.**
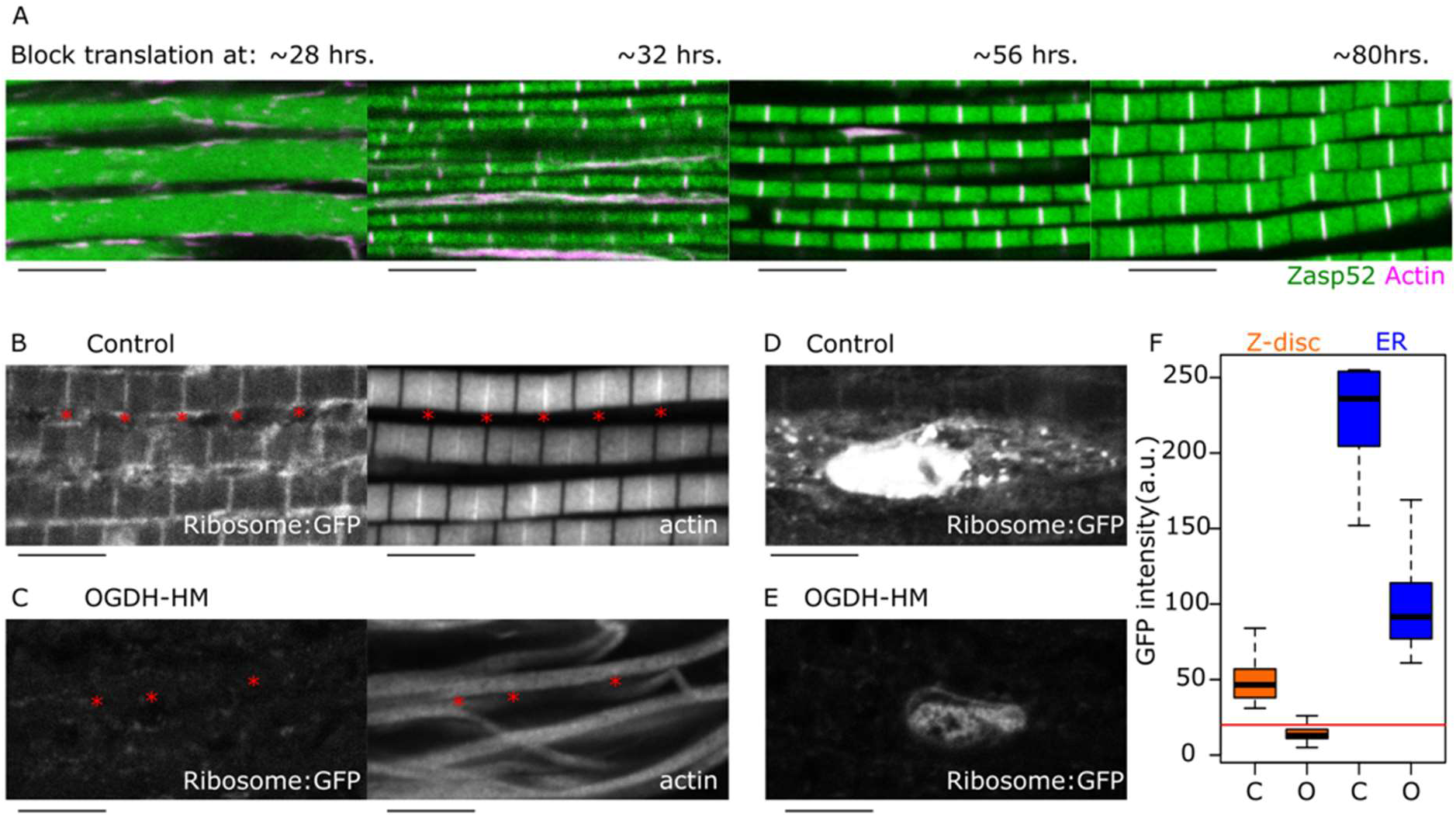
Amino acid balance is required for protein synthesis. (A) Blocking ribosome function using temperature-sensitive Ricin at different time points results in the differential blocking of myofibril diameter growth. Blocking ribosome function at ∼ 28 hours after pupa formation (APF) completely abolishes striation. Blocking ribosome function at ∼ 32 hours APF results in myofibril striation, but diameter growth is impaired. Blocking ribosome function at ∼ 56 hours APF results in smaller diameter but otherwise normal myofibrils. Blocking ribosome function at ∼ 32 hours APF does not affect myofibril appearance much. (B-C) Confocal images of control and OGDH-HM muscles expressing a GFP tagged version of RpS5a (Ribosome-GFP), actin filaments are also shown. (B) In control samples (Act88F-Gal4, UAS-RpS5a -GFP), ribosomes are spread in the cytoplasm but accumulate at the Z-disc. In OGDH-HM samples (Act88F-Gal4, UAS-RpS5a -GFP, UAS-OGDH-HM), Ribosomes are mostly absent. Red asterisks in B and C indicate selected Z-discs. (D) In controls, ribosome-GFP is also detected in the region surrounding the nucleus. (E) In OGDH-HM, ribosome-GFP is present in the ER. The scale bar is 5 µm. (F) Box plot of ribosome-GFP intensities in Z-disc and ER in control (C) and OGDH-HM (O) muscles. The red line marks the noise level.

## Discussion

We found that the oxoglutarate dehydrogenase enzyme is a Z-disc component required for growth of myofibrils. Our data suggest that OGDH is recruited to the Z-disc by the LIM2 domain of Zasp52 as part of the myofibril growth process. During development and growth of the Z-discs, OGDH functions together with the other TCA cycle enzymes to maintain a proper supply of amino acids. Without tight control of the amino acid pool, protein synthesis is blocked by degradation of ribosomes or by stopping ribosome assembly. Because the myofibril growth process requires an enormous number of proteins to be synthesized, a transient block of protein synthesis leads to myofibrils with growth defects. This is a novel mechanism that links amino acid homeostasis with the regulation of myofibril size control dependent on proteins of the Zasp/Enigma family (Gonzalez-Morales, Marsh, et al., 2019; Gonzalez-Morales, Xiao, et al., 2019).

### Unexpected localization of the OGDH complex at the Z-disc

Most TCA cycle enzymes localize to the mitochondria, but some have both mitochondrial and cytoplasmic forms (Chen et al., 2015). We show that most TCA cycle enzymes are required for myofibril growth and assembly. Because all the subunits of the OGDH complex are located at the Z-disc, the conversion of oxoglutarate into succinyl-CoA happens at the Z-disc. While we do not have localization data for the other TCA enzymes, we showed that their activity is required for myofibril development. A likely scenario is that only the OGDH complex functions at the Z-disc and the remaining enzymes are either mitochondrial or cytoplasmic.

Though metabolic enzymes are not typically recognized as myofibril components, their presence at the Z-disc is not entirely unprecedented. Six glycolytic enzymes that catalyze consecutive reactions along the glycolytic pathway localize to the Z-disc (Sullivan et al., 2003; Wojtas, Slepecky, von Kalm, & Sullivan, 1997). Our data together with previous data on glycolytic enzymes suggest that the Z-disc may be a common space for metabolic reactions that take place at the cytoplasm of muscles. Similarly, an interactome study using titin-BioID knock-in mice showed the presence of a number of enzymes at the Z-disc (Rudolph et al., 2020). Overall, our work demonstrates that the OGDH complex localizes to the Z-disc. At this location it appears to function as part of the TCA cycle to supply the amino acids needed for protein synthesis to promote myofibril growth.

Using dSTORM super resolution imaging, we show that the three subunits of the OGDH complex localize at the Z-disc and assemble into an asymmetric configuration in which the E3 subunit lies at the periphery of the complex. This is consistent with asymmetries observed by cryo-EM models of native α-keto acid dehydrogenase complexes from the *Chaetomium thermophilum* fungus (Kyrilis et al., 2021). Our work shows for the first time the asymmetry of the OGDH complex in vivo.

### OGDH is a novel Zasp-binding protein

Zasp proteins exist in two isoforms, the ones that contain LIM domains were designated as growing isoforms, while the ones that do not are called blocking isoforms (Katzemich et al., 2013; Liao, González-Morales, & Schöck, 2016). The growing isoforms recruit both growing and blocking forms through their LIM domains. The LIM domains bind the ZM domain present in both forms (Gonzalez-Morales, Xiao, et al., 2019). Here we show that the LIM2 domain of Zasp52 recognizes the OGDH protein through a sequence very similar to the ZM domain. Interestingly, while all LIM domains can bind Zasp66, only the LIM2 domain binds OGDH, indicating that the other LIM domains might be used exclusively for Zasp proteins or for yet unidentified Z-disc proteins. Screening for proteins with ZM-like regions might be a strategy to find other Zasp-binding proteins.

### Metabolic control of myofibril growth

Our data suggest a model in which OGDH with the TCA cycle maintain amino acid homeostasis, required to sustain the fast growth of myofibrils. When OGDH function is impaired, many amino acids appear to be affected, some accumulate, and others have highly reduced levels. The most affected amino acid is histidine which becomes practically undetectable. This is relevant because previous research has linked Histidine availability to ribosome biogenesis. In fast growing epithelial cells, high levels of Histidine are required for ribosome biogenesis (Froldi et al., 2019). In addition, the loss of SLC15, a histidine amino acid transporter, correlates with low cytoplasmic Histidine levels and with disruption of the mTOR pathway, which in turn controls ribosome biogenesis (Iadevaia et al., 2014; Kobayashi et al., 2014) While we have not explored the requirements of specific amino acids, we speculate that very low levels of Histidine, like the ones in OGDH-HM muscles, are sufficient to block ribosome biogenesis.

Several pathways are involved in coordinating *Drosophila* IFM growth. As sarcomere homogeneity is critical for muscle function, several pathways coordinate the growth process. At the global muscle level, the Hippo pathway controls the rapid growth burst of the flight muscles by controlling the expression of many of the major sarcomere proteins (Kaya-Copur et al., 2021), The activating function of RBf with the E2F/DP heterodimeric transcription factor promotes the postmitotic expression of sarcomere proteins required for myofibril growth (Zappia & Frolov, 2016; Zappia et al., 2019), and the Insulin and mTOR signaling pathways regulate endoreplication, required for postmitotic growth (Demontis & Perrimon, 2009). Then, at the local myofibril level, the Zasp oligomerization pathway controls myofibril diameter (Gonzalez-Morales, Xiao, et al., 2019) and a combination of titin filaments and actin regulators set the sarcomere length (Molnar et al., 2014; Shwartz et al., 2016; Tskhovrebova & Trinick, 2017). Finally, myofibril proteins are translated locally at the Z-disc (Denes, Kelley, & Wang, 2021). Here we add a link between the Zasp oligomerization system and the OGDH complex, which lies at the core of the TCA cycle. At the Z-disc, the OGDH complex directly impacts the local availability of amino acids, which in turn limits the rate of protein synthesis. Our data suggest a feedback system between Zasp oligomerization and the local amino acid pool that provides robustness to the myofibril diameter size control.

## Methods

### Experimental model

As a model organism, we used *Drosophila melanogaster*. Flies were raised at 25°C. A comprehensive list of all strains used and generated can be found in the Key Resources Table. We used Saccharomyces cerevisiae for the yeast two-hybrid assays. The UAS/Gal4 system was used for transgene expression. The Act88F-Gal4 transgene was used to direct expression in the indirect flight muscles (Bryantsev et al., 2012).

### Confocal microscopy imaging of flight muscles

The muscles were prepared for confocal imaging as described previously (Xiao et al., 2017). Briefly, the thoraces were dissected in halves and incubated overnight at -20°C in Relaxing-Glycerol solution (20 mM Na-Phosphate pH 7.2, 2 mM MgCl2, 2 mM EGTA, 5 mM DTT, 0.5% Triton X-100, 50% glycerol). Then, we fixed the muscles in 4% paraformaldehyde and dissected the muscles. For visualizing actin filaments, we used 488-phalloidin or 555-phalloidin (1:1000; Cytoskeleton) in PBS. Finally, we mounted the samples in Mowiol 4-88 mounting media (Sigma 9002-89-5). All images were acquired using a 63x 1.4 NA HC Plan Apochromat oil objective on a Leica SP8 confocal microscope. We used more than 10 flies for each experiment and randomly picked the muscle area to image. Control and experimental samples were prepared and imaged simultaneously and imaged with comparable parameters.

### Bimolecular fluorescence complementation assay

BiFC assays were done as previously described (Marescal et al., 2020). The UAS-OGDH-NYFP construct was made using Gateway cloning using the OGDH-GEO09867 donor vector that contains the PA isoform as donor and pBIDUAS-GV, pUAST-RfB-myc-NYFP as destination vector (Gohl, Banovic, Grevelhorster, & Bogdan, 2010). To make the UAS-OGDH-BM-NYFP construct, we first deleted the coding sequence for amino acids 741-769 deletion into the OGDH-GEO09867 donor vector. Then the resulting vector was transferred to pUAST-RfB-myc-NYFP using Gateway cloning. The resulting vectors were sequence verified and then inserted into the ZH-58A attp landing site. The UAS-Zasp52-PK-CYFP, and the control lines were described previously (Gohl et al., 2010; Gonzalez-Morales, Xiao, et al., 2019). At least 10 samples were used for each condition. We normalized the data to the basal noise levels and made plots in R software.

### Tissue-specific CRISPR mutants

Tissue-specific CRISPR disruption works by expressing the Cas9 endonuclease in a specific tissue, using the UAS/Gal4 system together with a gene targeting gRNA expressed ubiquitously. The tissue containing the Cas9 protein generates small insertion or deletion mutations in the gene targeted by the gRNA (Port, Chen, Lee, & Bullock, 2014) To express the Cas9 protein in the IFM, we used Act88F-Gal4 with UAS-Cas9.P2 (BDSC # 58986). As gRNA constructs, we used TKO.GS03432 targeting CG5214/DLST, TKO.GS00548 targeting CG7430/DLD, and TKO.GS00550 targeting Nc73EF/OGDH.

### Construction of DLST-GFP line

The fosmid carrying the GFP tagged version of DLST (SourceBioscience: CBGtg9060A03104D) is part of the flyfos library, a collection of fosmids that contain C-terminally GFP tagged versions of genes at their genomic locations (Sarov et al., 2016). These constructs are then introduced into the fly genome by site-directed integration using the PhiC31 integrase (Bischof, Maeda, Hediger, Karch, & Basler, 2007). We used P[CaryP]attP40 as the landing site for the DLST-GFP fosmid. Genome ProLab did the microinjections and Px3-RFP was used to screen for successful transformants.

### Precise genome engineering at the OGDH locus

To precisely modify the OGDH/Nc73EF locus, we used the Recombination-Mediated Cassette Exchange method using *OGDH*^*MI06026*^, which carries a MiMIC transposon between exons 5 and 6 (Venken et al., 2011). The rationale was to replace the MiMIC transposon with wild type or mutant versions of the OGDH gene starting with the sequence where *OGDH*^*MI06026*^ is inserted. In all cases, we added a C-terminal tag consisting of 6XHis and mCherry. First, we gene synthesized the wild-type replacement construct and then mutagenized that construct using site-directed mutagenesis in bacteria. Gene synthesis and mutagenesis was done by Genscript. We did *OGDH*^*H306A-H344A*^ (*OGDH-ED*), an enzymatic dead version of OGDH, by replacing the conserved H306 and H344 with alanines. These residues are required for oxoglutarate binding (Frank et al., 2007). We did *OGDH*^*Δ741-769*^ (*OGDH-BM*) by deleting residues 741 to 769. Residue numbering is done according to the OGDH-PA isoform. All constructs were then introduced into *OGDH*^*MI06026*^ as a landing site. GenetiVision did the microinjections and initial confirmation of the mutants.

### Yeast two-hybrid assays

Yeast two-hybrid assays were done as described previously (González-Morales, Xiao, et al., 2019).

### GC/MS Sample Preparation and Metabolite Measurements

The thoraces were dissected then flash frozen and crushed by mortar and pestle on liquid nitrogen (40 thoraces per sample). Frozen tissue powder was placed in pre-chilled Eppendorf brand tubes to which 1 ml of 80% methanol/water was added along with four 2.8 mm ceramic beads. Samples were subjected to 45 sec of bead beating at 50 Hz (SpeedMill Plus homogenizer) 4 times. Samples were kept on ice between bead beating sessions. Samples were then centrifuged at 1°C for 10 min and 15,000 rpm (21,130 rcf). Supernatants were transferred to fresh pre-chilled tubes containing 1 µl of 800 ng/µl 2H27-Myristic in pyridine. The protein concentration of the pellets was estimated and used for normalization. Samples were then dried by vacuum centrifugation operating at a sample temperature of -4°C (LabConco).

After drying, samples were subjected to a two-step derivatization: First the samples were resuspended in 30 µl of 10 mg/ml Methoxyamine:HCl in anhydrous pyridine (MOX). They were sonicated and vortexed 15 seconds three times then centrifuged for 3 min at room temperature and 15,000 rpm (21,130 rcf). Incubation for methoximation was 30 min at room temperature. The samples were then centrifuged for 2 min at 15000 rpm (21,130 rcf) and the supernatants were transferred to GC/MS sample vials containing 250 µl glass inserts pre-filled with 70 µl of N-tert-butyldimethylsilyl-N-methyltrifluoroacetamide (MTBSTFA) and incubated at 70°C for 60 min.

An Agilent 5975C GC-MS equipped with a DB-5MS+DG (30 m x 250 µm x 0.25 µm) capillary column (Agilent J&W, Santa Clara, CA, USA) was used for all GC-MS measurements, and data collected by electron impact set at 70 eV both in scan (50-1000 m/z) and single ion monitoring modes. A volume of 1 mL of derivatized sample was injected in splitless mode with inlet temperature set to 280°C, using helium as a carrier gas and the flow rate adjusted to 18 min for 2H27-myristic acid. The quadrupole was set at 150°C and the GC/MS interface at 285 °C. The oven program for all metabolite analyses started at 60°C held for 1 min, then increasing at a rate of 10°C/min until 320 °C. Bake-out was at 320°C for 10 min. Sample data were acquired in scan mode (50-1000 m/z) or in single ion monitoring (SIM) with a 5 ms dwell time where the M-57 [M+·- C4H9·]+ fragment was used for quantitation (area under the curve) in both modes of data acquisition. Citrate and isocitrate used the M/z 459 ion for quantification as described previously (Mamer et al., 2013). The spectra and retention times of all metabolites reported were confirmed by methoxylamine – tert-butyldimethylsilylated authentic standards. For saturating metabolites, samples were diluted 1:25 with the same ratio of derivatization reagents and ran in scan mode. Metabolite area under the curve was normalized to tissue weight.

### Transmission electron microscopy

Samples were prepared for transmission electron microscopy imaging as described previously with slight modifications (González-Morales et al., 2017). Briefly, the thoraces were dissected in halves and were treated with 5 mM MOPS pH 6.8, 150 mM KCl, 5 mM EGTA, 5 mM ATP, 1% Triton X-100 for 2 hours at 4°C. Samples were then washed in rigor solution (5 mM MOPS pH 6.8, 40 mM KCl, 5 mM EGTA, 5 mM MgCl2, 5 mM NaN3) and fixed in 3% glutaraldehyde, 0.2% tannic acid in 20 mM MOPS pH 6.8, 5 mM EGTA, 5mM MgCl2, 5 mM NaN3 for 2 hours at 4°C. Images were acquired on a Tecnai 12 BioTwin 120 kV transmission electron microscope with an AMT XR80C CCD camera (FEI).

### Super resolution dSTORM microscopy

Super resolution imaging was done essentially as described previously (Szikora et al., 2020). Briefly, all the dSTORM images were captured under EPI illumination (Nikon CFI Apo 100x, NA=1.49) on a custom-made inverted microscope based on a Nikon Eclipse Ti-E frame. The laser (MPB Communications Inc.: 647 nm, Pmax=300 mW) intensity controlled via an acousto-optic tunable filter (AOTF) was set to 2-4 kW/cm2 on the sample plane. An additional laser (Nichia: 405 nm, Pmax=60 mW) was used for reactivation. Images were captured by an Andor iXon3 897 BV EMCCD digital camera (512×512 pixels with 16 μm pixel size). Frame stacks for dSTORM super-resolution imaging were captured at a reduced image size. A fluorescence filter set (Semrock, LF405/488/561/635-A-000) with an additional emission filter (AHF, 690/70 H Bandpass) were used to select and separate the excitation and emission lights in the microscope. During the measurements, the perfect focus system of the microscope was used to keep the sample in focus with a precision of <30 nm. Right before the measurement the storage buffer of the sample was replaced with a GLOX switching buffer (van de Linde et al., 2011) and the sample was mounted onto a microscope slide. Typically, 20,000-50,000 frames were captured with an exposure time of 20 or 30 ms. The captured and stored image stacks were evaluated and analyzed with the rainSTORM localization software (Rees, Erdelyi, Schierle, Knight, & Kaminski, 2013). Individual images of single molecules were fitted with a Gaussian point spread function and their center positions were associated with the position of the fluorescent molecule. Localizations were filtered via their intensity, precision, and standard deviation values. Only localizations with precisions of <20 nm and standard deviation between 0.8≤σ≤1.0 were used to form the final image and for further analysis. Mechanical drift introduced by either the mechanical movement of the sample or thermal effects was analyzed and reduced by means of a correlation based blind drift correction algorithm. Spatial coordinates of the localized events were stored, and the final super-resolved image was visualized with a pixel size of 10 nm.

### Ribosome block

To block protein synthesis, we used UAS-RA.cs2, a cold-sensitive version of the Ricin-A toxin subunit. Ricin-A inactivates ribosomes by the specific depurination of the 28S rRNA (Moffat et al., 1992). Ricin-A-TS is a temperature sensitive allele that is active at 30°C but not at 20°C. We raised Act88F-Gal4 UAS-Ricin-A-TS flies at 20°C then transferred them to a 30°C incubator for 48 hours, and then back into 20°C. The muscles were analyzed 2 days after emergence.

## Acknowledgments

We appreciate the help of Beili Hu in making transgenic flies. We appreciate the support from the community resources like the Bloomington Drosophila Stock Center, Fybase, and the Vienna Drosophila Resource Center. Confocal imaging was done at the Advanced Bioimaging Facility. TEM was done with the assistance of Jeannie Mui at the Facility for Electron Microscopy Research. All GC/MS data were collected at the Metabolomics Innovation Resource. This work was supported by operating grants MOP-142475 and PJT-155995 from the Canadian Institutes of Health Research.

## Figure Legends

**Figure 1—figure supplement 1. Additional characterization of OGDH-HM myofibril phenotype**. (A) TEM image of an aggregate formed in OGDH-HM muscles. Notice the actin filaments that connect to the aggregate. Scale bar is 1 µm. (B-E) Confocal images of OGDH-HM muscles with different sarcomere proteins labelled. In all cases Zasp52 marks the Z-discs and the aggregates in magenta. Scale bars are 5 µm. (B) Actinin localizes to the small Z-discs and the aggregates. (C) Zasp66 localizes to the small Z-discs and the aggregates. (D) Sls/Titin localizes to the Z-discs but not the aggregates. (E) The M-line protein Obscurin does not localize to the Z-discs or the aggregates. (F-H) Confocal images of muscles in three alternative approaches to deplete OGDH. Actin filaments are shown in green and Zasp52 marks the Z-discs in magenta. (F) The RNAi GD12778 directed against OGDH results in small aggregates and myofibril defects. (G) The RNAi GD50393 directed against another region of OGDH results in myofibril disintegration. (H) The TKO CRISPR-based method directed against OGDH results in complete disintegration of myofibril structure. (I) Boxplot of Z-disc heights in different OGDH depleted conditions. (J) Plot of the surface area of Zasp52 positive particles. Small particles are Z-discs while large particles are aggregates. Notice the large number of aggregates in OGDH depleted conditions.

**Figure 4—figure supplement 1** (A) Reference I-band image showing the distribution patterns of Actinin, MyosinS2 and Zasp52 epitopes. (B) Longitudinal epitope average normalized localization epitope densities of the three OGDH complex subunits relative to the distribution of Myosin S2 epitope distribution pattern.

**Figure 4—figure supplement 2** Details of the dSTORM super resolution imaging experiment. An example of a widefield image, examples of individual dSTORM images, the average distribution image, and descriptive statistics of the distribution patterns are shown for each subunit of the OGDH complex. The scale bar is 500 nm. The number of individual dSTORM images is shown in the average image. In the average dSTROM image, n represents the number of individual dSTORM images used to create the distribution average image. In the statistics, n represents the number of individual dSTROM average images.

**Figure 5—figure supplement 1. ATP levels in different TCA cycle enzyme deficiencies**. Bar plot of the relative ATP levels to the control are shown for 8 enzymes of the TCA cycle and for 2 Cox5A mutant conditions. As expected, reducing Cox5A greatly affects ATP production while OGDH depletion has no effect on reducing ATP levels. p-Values were calculated using Welch’s two-sample t-test.

