## Supplementary figures and images for "Oxoglutarate dehydrogenase coordinates myofibril growth by maintaining amino acid homeostasis"

### Figure 1 - figure supplement 1

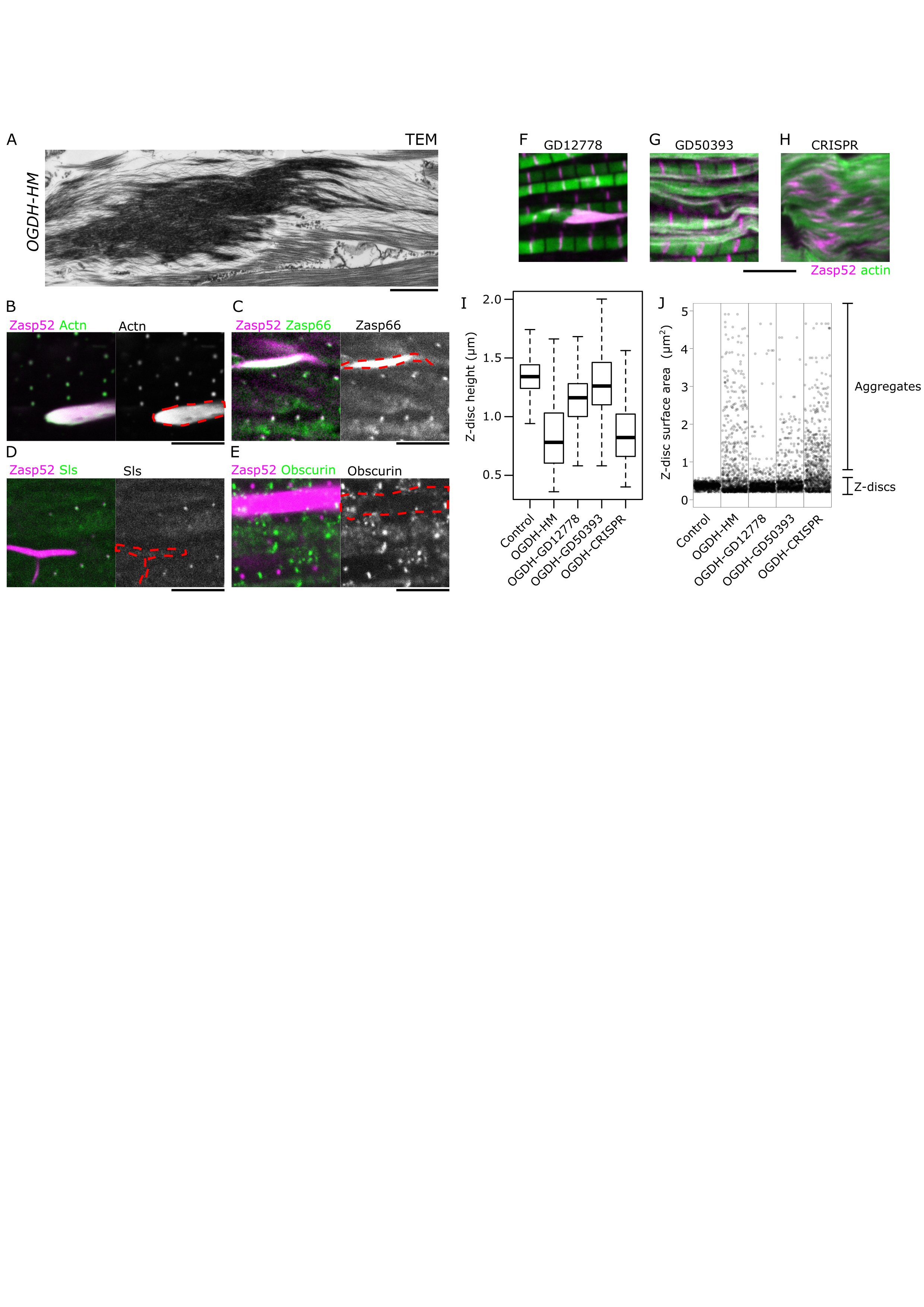

### Figure 4 - figure supplement 1

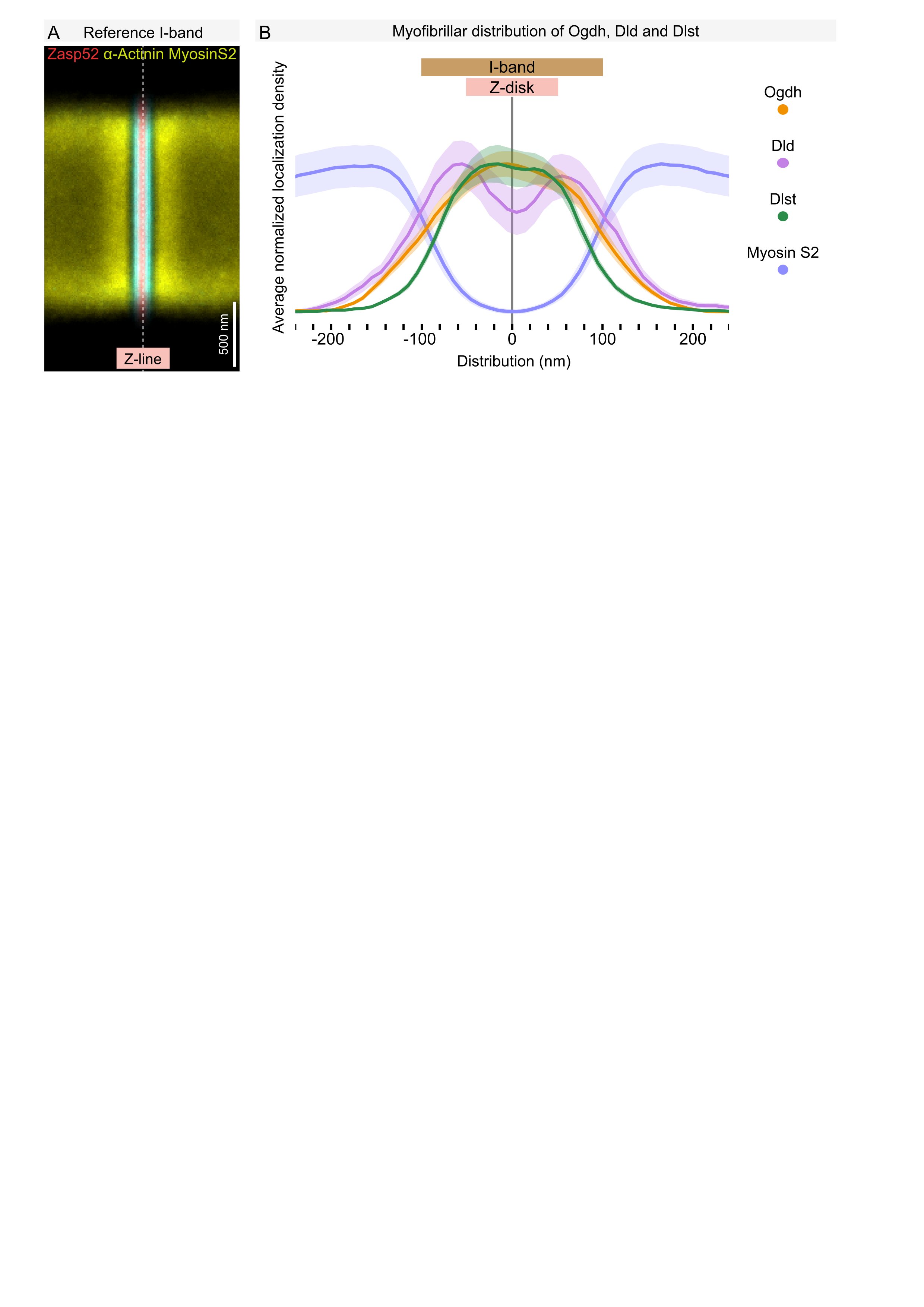

### Figure 4 - figure supplement 2

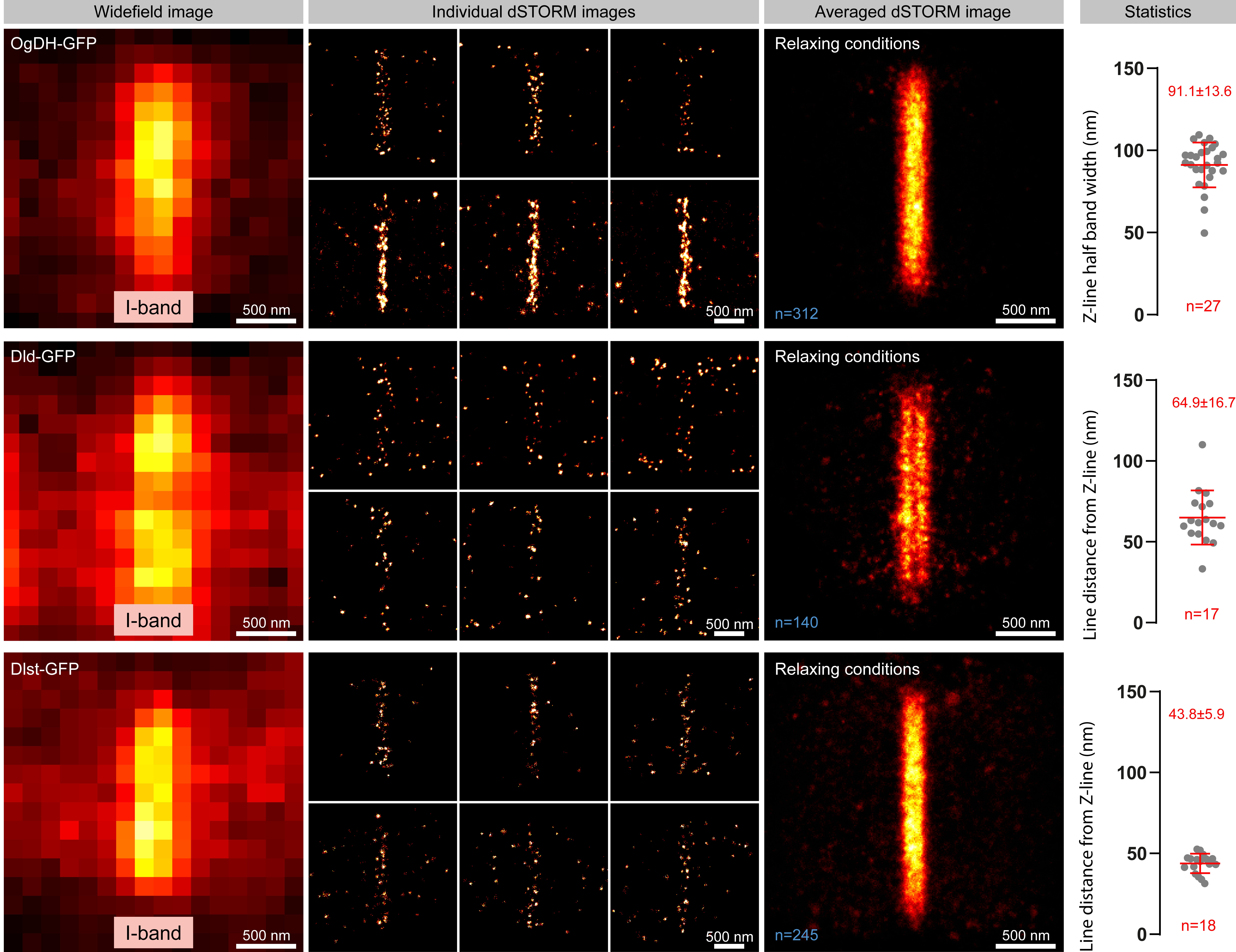

### Figure 5 - figure supplement 1

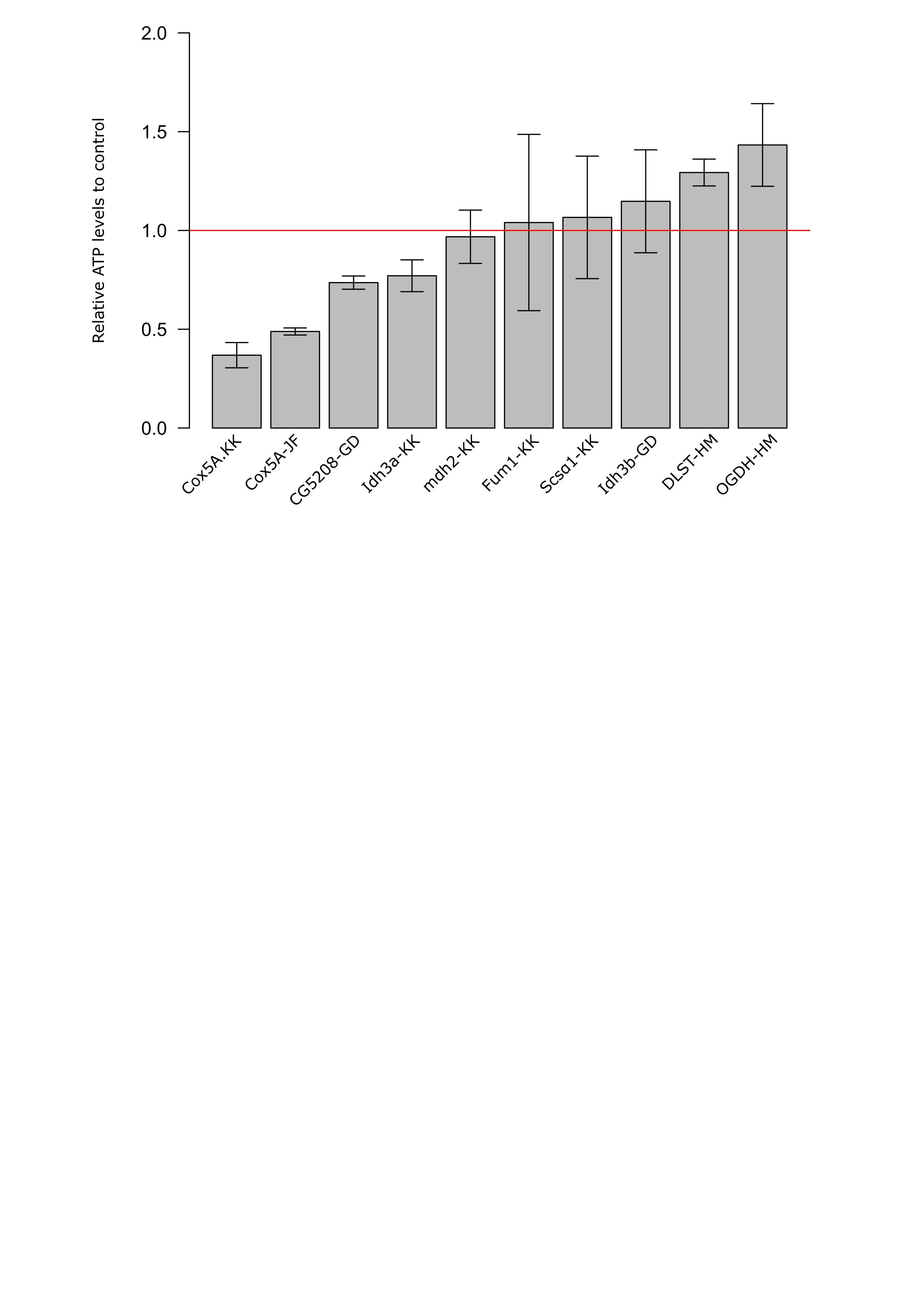
